# Neural discrimination of emotional prosody in natural sentences in 2- to 3-month-old infants using OPM-MEG

**DOI:** 10.64898/2026.11.03.763724

**Authors:** Jaeyoung Yang, Valeria Monterrubio Leija, Alice Waitt, Anna Kowalczyk, Andrew Bremner, Barbara Pomiechowska, Giulia Orioli, Ole Jensen, Kyung-min An

**Affiliations:** Centre for Human Brain Health, School of Psychology, University of Birmingham, Birmingham B15 2TT, UK; Centre for Developmental Science, School of Psychology, University of Birmingham, Birmingham B15 2TT, UK; School of Computer Science, University of Birmingham, Birmingham B15 2TT, UK; College of Medicine and Health, University of Birmingham, Birmingham B15 2TT, UK; Department of Experimental Psychology, University of Oxford, Oxford, UK; Oxford Centre for Human Brain Activity, Oxford Centre for Integrative Neuroimaging, Department of Psychiatry, University of Oxford, Oxford, UK

**Keywords:** Emotional prosody, Infants, OPM-MEG, Multivariate pattern analysis, Neural decoding, Speech perception

## Abstract

From birth, infants are sensitive to the emotional prosody of speech, yet it remains unclear when the developing brain discriminates emotions conveyed by natural spoken sentences. Here, we recorded brain activity with optically pumped magnetometer magnetoencephalography (OPM-MEG) in 15 infants aged 2-3 months while they listened to semantically neutral sentences spoken with happy or angry prosody. Natural sentences elicited robust auditory evoked responses, with prominent components at 88, 216, 380 and 1,356 ms after sentence onset and sources centred on bilateral superior and middle temporal cortex. Time-resolved multivariate pattern analysis revealed that distributed neural activity discriminated recordings spoken with happy from angry prosody 346-454 ms after the onset of the prosodically prominent word (the first word carrying a pitch peak), despite no difference in global field power between conditions. This discrimination was transient, emerged before the end of the sentence, and was carried primarily by phase-locked delta-band activity, followed by later differences in induced alpha power. Angry prosody also delayed the late auditory response by approximately 15 ms with no detectable changes in amplitude. Our findings show that cortical responses to happy and angry prosody in natural speech differ by 2-3 months of age, and demonstrate that OPM-MEG combined with time-resolved decoding can reveal the temporal, spatial and spectral organisation of early emotional-prosody processing.

## Introduction

Humans are particularly sensitive to others’ voices from birth, and the voice, together with facial expressions, gaze and tactile interaction, is a key channel for forming early social relationships (DeCasper & Fifer, 1980; Grossmann, 2010a; Gibson & Pick, 2000). Speech prosody, that is, variation in pitch, rhythm, stress and speech rate, can convey a speaker’s emotional state even before the meaning of a sentence is fully understood (Banse & Scherer, 1996), and is likely to be a primary cue from which young infants obtain emotional information (Zhang et al., 2019). Because infants must acquire information about threat, safety and attachment from their caregivers’ voices even when visual cues are limited, early sensitivity to emotional prosody may form a foundation for social and emotional development (Vaish & Striano, 2004; Vaish et al., 2008).

Previous studies have shown that infants are sensitive to emotional prosody, and that this sensitivity is present from shortly after birth. Newborns open their eyes more when hearing happy speech in their native language (Mastropieri & Turkewitz, 1999), show different sucking responses to happy sentences than to sad and neutral sentences (Silvestri et al., 2026). Differences in brain responses to emotional prosody have also been reported. With electrophysiological measures, newborns showed EEG mismatch responses to emotional syllables (Cheng et al., 2012) and 3- to 12-month-old infants showed mismatch responses to emotional prosody in spoken words (Kao & Zhang, 2025), and 7-month-old infants showed distinct ERPs to emotional words (Grossmann et al., 2005). With haemodynamic measures, responses in temporal, frontal and parietal regions differed with emotional prosody in fNIRS studies of newborns (Zhang et al., 2017, 2019), a diffuse optical tomography study of 2-month-old infants (Shekhar et al., 2019) and an fNIRS study of 7-month-old infants (Grossmann et al., 2010b), and an fMRI study of 3- to 7-month-old infants found that sad non-speech vocalisations modulated activity in the orbitofrontal cortex and insular (Blasi et al., 2011). However, EEG offers millisecond resolution but limited spatial precision, whereas haemodynamic methods such as fNIRS, diffuse optical tomography and fMRI provide spatial information but cannot resolve fast neural dynamics because the blood-flow response is slow. It therefore remains unclear how the processing of emotional prosody is organised in time, space and frequency during the first months of life, when the auditory cortex is still immature. Addressing this question requires a method with both high temporal and high spatial resolution.

Optically pumped magnetometer-based magnetoencephalography (OPM-MEG) has millisecond temporal resolution like EEG, far higher than that of fNIRS and fMRI, and offers higher spatial resolution than EEG, because it measures magnetic fields rather than electrical potentials, which are strongly distorted as they pass through the scalp and skull (Hämäläinen et al., 1993; Baillet, 2017). OPM-MEG sensors are smaller and lighter than conventional helium-based SQUID-MEG sensors and can be positioned flexibly, so helmets can be built to fit the size and shape of infants’ heads, opening new possibilities for infant research (Boto et al., 2018; Hill et al., 2019; Brookes et al., 2022; Tierney et al., 2019). Because the sensors can be placed close to the scalp, source–sensor distances are short, yielding larger signals even from infants’ small heads (Tierney et al., 2019; Brookes et al., 2022). Auditory responses have recently been recorded with OPM-MEG in 1-month-old (Corvilain et al., 2025) and 2- to 3-month-old infants (Orioli et al., 2026). In 1-month-old infants, auditory response amplitudes measured with OPM-MEG were about four times larger than SQUID-MEG responses in the same infants (Corvilain et al., 2025). In the present study, the high spatiotemporal resolution of OPM-MEG can therefore reveal two aspects of early emotional prosody: when emotional information appears within infant auditory responses, that is, the time course of emotional responses early in life, and in which brain regions this processing is organized.

The brain represents information in patterns of activity distributed across multiple regions rather than in the magnitude of activity in a single region (Haxby et al., 2001; Kriegeskorte et al., 2006). Time-resolved multivariate pattern analysis (MVPA) is a machine-learning approach that exploits such patterns. At each time point, a classifier is trained on signals from multiple sensors, and the information carried by neural activity at that time point is quantified by how accurately the classifier predicts the condition in data not used for training (Carlson et al., 2013; King & Dehaene, 2014). This approach can read out condition information from patterns distributed across sensors even when the global response magnitude does not differ between conditions (Grootswagers et al., 2017; Cichy & Pantazis, 2017), and it has been applied to infant data with few trials and weak signals (Ashton et al., 2022; Bayet et al., 2020). Moreover, transforming classifier weights into activation patterns makes it possible to identify which sensors and brain regions carry the information used by the decoder (Haufe et al., 2014). Because the information used for decoding ultimately lies in spatial patterns across sensors, OPM-MEG is expected to provide the decoder with finer spatiotemporal patterns. Combining OPM-MEG with MVPA should therefore make it possible to exploit the spatial organisation of infant responses to track, at millisecond resolution, when emotional information is present and where it arises.

In everyday life, the emotional prosody that infants hear is embedded not in isolated syllables but in continuous speech, and the pitch contours and rhythms that distinguish emotions unfold across multiple syllables and words (Banse & Scherer, 1996). However, previous infant studies of emotional prosody with millisecond temporal resolution have mainly used single syllables or words, and few have examined at the millisecond scale how happy and angry prosody unfolds over time within natural sentences (Grossmann et al., 2005; Cheng et al., 2012; Kao & Zhang, 2025). By contrast, studies of sentence-level prosody have relied on behavioural and haemodynamic measures (Mastropieri & Turkewitz, 1999; Zhang et al., 2017, 2019; Shekhar et al., 2019; Silvestri et al., 2026), Consequently, the time course with which emotional prosody in natural sentences is processed in the infant brain remains unknown. In the present study, we therefore examine brain responses to emotional prosody within short semantically neutral spoken sentences lasting more than one second.

Using OPM-MEG and time-resolved MVPA, this study addresses three questions. First, what temporal structure do auditory responses to natural spoken sentences show in the brains of 2- to 3-month-old infants? Second, when and in which brain regions do happy and angry prosody give rise to discriminable neural patterns? Third, on which frequency bands, particularly delta-, theta- and alpha-band phase-locked activity or induced power, does this discrimination depend? Through these questions, this study examines how the sensitivity to emotional prosody reported from the neonatal period is expressed in temporal, spatial and spectral neural organisation during the first three months of life.

## Materials and Methods

### Participants

Seventeen healthy infants aged 2–3 months (52–99 days; mean ± SD: 73.0 ± 16.4 days; 9 females, 8 males) from English-speaking households were recruited through the Birmingham BabyLab Database. All infants were born at ≥37 weeks of gestation and had no history of neurological or medical conditions, hearing impairment, or implanted metallic devices. Accompanying caregivers had no implanted metallic devices or history of claustrophobia. Two infants were excluded because they contributed fewer than 50 artefact-free trials per condition. The final sample comprised 15 infants (71.7 ± 16.7 days; 8 females, 7 males; Table 1). The study was approved by the University of Birmingham STEM Ethics Committee (ERN_23-1163P). Caregivers received a full explanation of the study procedures and provided written informed consent before participation.

**Table 1.**
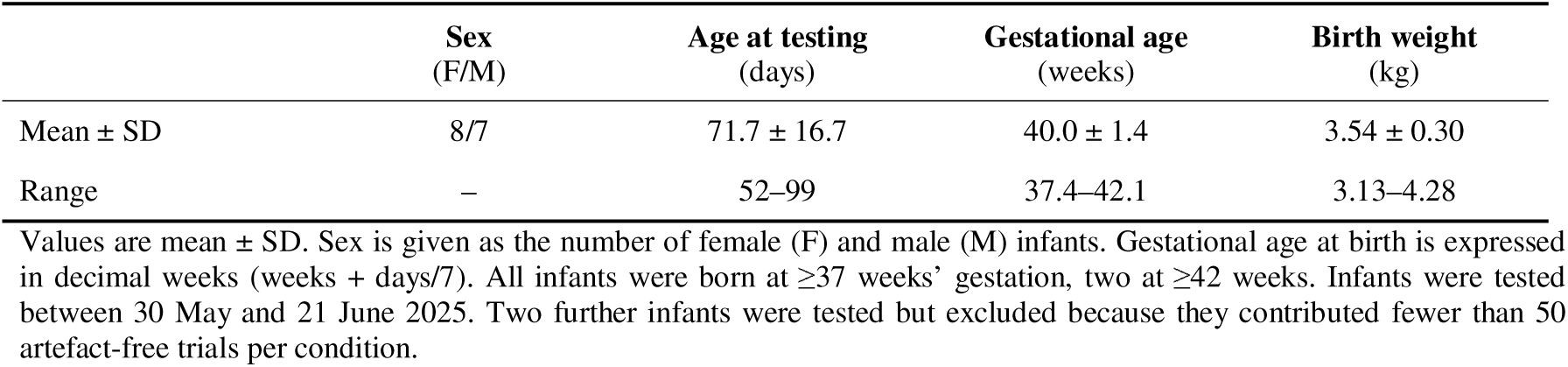
Participant characteristics (*n* = 15)

|  | **Sex**<br>(F/M) | **Age at testing**<br>(days) | **Gestational age**<br>(weeks) | **Birth weight**<br>(kg) |
|---|---|---|---|---|
| Mean ± SD | 8/7 | 71.7 ± 16.7 | 40.0 ± 1.4 | 3.54 ± 0.30 |
| Range | – | 52–99 | 37.4–42.1 | 3.13–4.28 |

Values are mean ± SD. Sex is given as the number of female (F) and male (M) infants. Gestational age at birth is expressed in decimal weeks (weeks + days/7). All infants were born at ≥37 weeks' gestation, two at ≥42 weeks. Infants were tested between 30 May and 21 June 2025. Two further infants were tested but excluded because they contributed fewer than 50 artefact-free trials per condition.

### Auditory stimuli

Auditory stimuli comprised semantically neutral sentences, such as “the book was green”, spoken by female British speakers with happy or angry prosody (Fig. 1A). The sentences therefore conveyed emotion through their vocal delivery without explicitly emotional semantic content. Recordings were selected from an existing emotional prosody corpus (Paulmann et al., 2016) and used with permission. Peak levels of the recordings ranged from −1.8 to −0.8 dBFS. To select readily recognisable exemplars, 22 adult listeners categorised each recording as happy, angry, or neutral. Recognition accuracy was calculated as the proportion of listeners whose responses matched the intended emotional category. The ten recordings with the highest adult recognition accuracy in each of the happy and angry categories were selected, yielding 20 stimuli.

**Figure 1.**
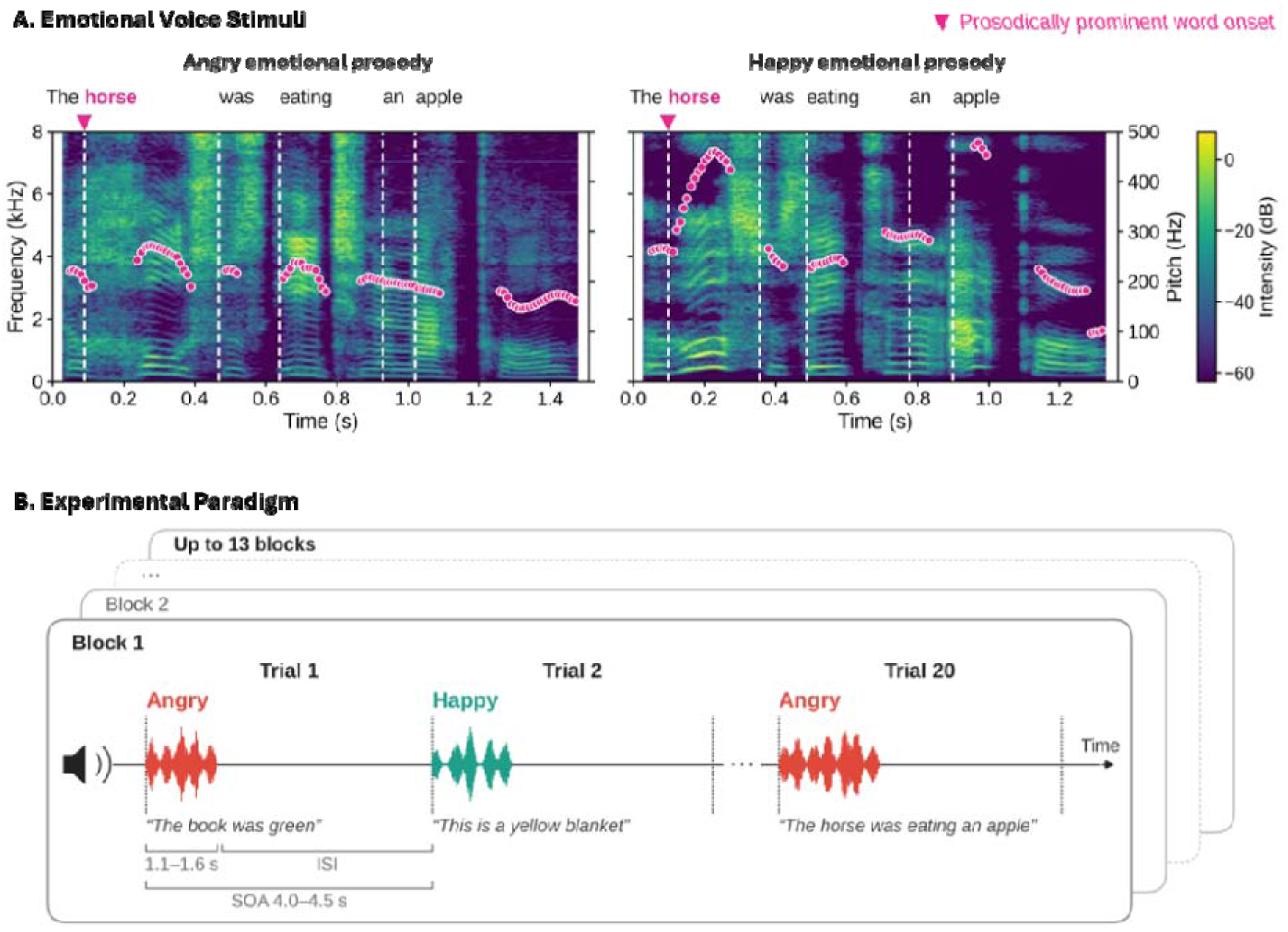
Auditory stimuli and experimental paradigm. **(A)** Spectrograms of an angry (left) and a happy (right) recording of “The horse was eating an apple”, plotted on the same time scale; colour indicates intensity (dB, scaled separately for each recording). Circles show the F0 contour (right axis), and white dashed lines mark word onsets. Time 0 is sentence onset; arrowheads mark the onset of the prosodically prominent word, the word containing the first detected F0 peak (“horse”). **(B)** Paradigm. Infants heard up to 13 blocks of 20 trials, each block containing the ten happy and ten angry recordings in pseudorandom order, with no more than two consecutive trials of the same emotion. Recordings lasted 1.07–1.63 s, and the stimulus onset asynchrony (SOA) varied randomly between 4.0 and 4.5 s.

Stimulus durations ranged from 1.07 to 1.63 s (1.37 ± 0.14 s). Happy and angry recordings did not differ in duration, number of words, or mean intensity, but mean F0 was lower for angry (218.0 ± 17.4 Hz) than for happy recordings (269.5 ± 28.5 Hz; *t*(18) = −4.88, *P* < 0.001). Stimulus characteristics are summarised in Table 2.

**Table 2.** Stimulus characteristics.

|  | **Happy**<br>(*n* = 10) | **Angry**<br>(*n* = 10) | ***t*(18)** | ***P*** |
|---|---|---|---|---|
| *Linguistic content* |  |  |  |  |
| Words per sentence | 5.3 ± 0.5 | 5.0 ± 0.7 | −1.15 | 0.26 |
| Syllables per sentence | 6.7 ± 0.7 | 6.1 ± 1.3 | −1.31 | 0.21 |
| *Acoustic properties* |  |  |  |  |
| Duration (s) | 1.34 ± 0.08 | 1.40 ± 0.18 | 1.05 | 0.31 |
| Mean F0 (Hz) | 269.5 ± 28.5 | 218.0 ± 17.4 | −4.88 | < 0.001* |
| Mean intensity (dB) | 68.2 ± 2.7 | 65.9 ± 4.1 | −1.48 | 0.16 |

Values are mean ± s.d. across recordings. *t* and *P* values are from two-sided independent-samples Student’s *t*-tests (df = 18) comparing angry with happy recordings; negative *t* values indicate lower values for angry recordings. Duration is the total file duration. Mean F0 was computed over voiced frames (Parselmouth/Praat defaults: 75–600 Hz, 10-ms time step; Jadoul et al., 2018), and mean intensity is the average of the intensity contour (dB).

### Acoustic feature extraction and temporal landmarks

Stimulus acoustics were characterised using Parselmouth, a Python interface to Praat (Jadoul et al., 2018). Spectrograms were generated after pre-emphasis using a 30-ms analysis window and a maximum analysis frequency of 8,000 Hz (Fig. 1A). F0 contours were extracted for each recording using Praat’s pitch-estimation procedure. Word boundaries were obtained using the Montreal Forced Aligner (MFA; McAuliffe et al., 2017). Candidate F0 peaks were identified using SciPy’s find_peaks function (Virtanen et al., 2020), with permitted peak-height ranges of 100–400 Hz for angry recordings and 100–490 Hz for happy recordings and a minimum separation of 20 F0-contour samples between peaks. In each recording, the first detected F0 peak was selected, and the word containing it was defined as the prosodically prominent word. The onset of this word, rather than the time of the F0 peak itself, served as the primary reference point for aligning the neural responses, and sentence onset served as a secondary reference point (Fig. 1A). The prosodically prominent word began 57 ± 32 ms after sentence onset (range 0–100 ms; it was the first word in 4 of the 20 recordings), and its onset did not differ between conditions (angry 60 ± 23 ms, happy 54 ± 40 ms; *t*(18) = 0.41, *P* = 0.68). Utterance offsets occurred 1.33 ± 0.15 s after sentence onset (range 1.04–1.60 s) and did not differ between conditions (*t*(18) = 1.51, *P* = 0.15).

### Experimental paradigm

Stimuli were presented using Psychtoolbox (Brainard, 1997; Kleiner et al., 2007) running in MATLAB R2019b and delivered through a DATAPixx and SoundPixx system (VPixx Technologies, Canada) connected to a piezoelectric speaker (SPS series; Sonitron) positioned in front of the bed at approximately 2 m above the floor. Stimuli were presented at approximately 65 dB at the infant’s head position. Stimulus onsets were marked by digital triggers sent from the DATAPixx to the OPM-MEG acquisition system. The planned paradigm comprised 13 blocks of 20 trials, with each block containing the ten happy and ten angry recordings. Stimuli were presented in a pseudorandom order, with no more than two consecutive trials from the same emotional category. The stimulus onset asynchrony varied randomly between 4.0 and 4.5 s (Fig. 1B). When a run was interrupted, stimulation continued in a subsequent run once the infant had resettled. The number of trials presented thus varied across infants (happy: 117.7 ± 9.2, range 96–130; angry: 117.9 ± 9.5, range 94–130) but did not differ between conditions (*t*(14) = −0.72, *P* = 0.49). The total recording duration was 20.2 ± 2.4 min (range: 15.6–25.9 min), including breaks as needed by the infants, and the total visit, including preparation, lasted approximately 60–90 min.

### Infant OPM-MEG recording

Brain activity was recorded with 50 optically pumped magnetometer (OPM) sensors (HEDscan; FieldLine Inc., Boulder, CO, USA) in a rigid infant helmet (Fig. 2A), 3D-printed from SLA resin (circumference 45 cm) for infants aged 2–3 months; 50 of its 96 sensor slots were populated to provide whole-head coverage. The helmet was mounted at the head end of a purpose-built infant bed (Fig. 2B; Orioli et al., 2026), whose sliding platform allowed stable positioning of the infant’s head within the sensor array (Fig. 2C).

**Figure 2.**
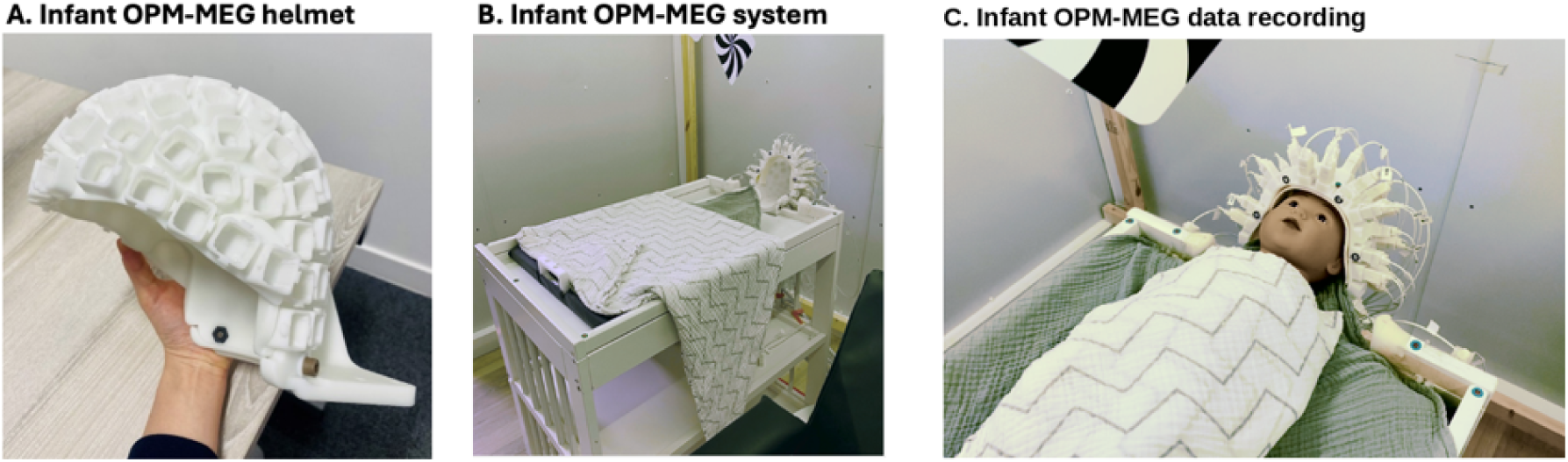
Infant OPM-MEG recording setup. **(A)** The 3D-printed helmet (FieldLine Inc.), designed for infants aged 2–3 months, shown before the sensors were fitted. **(B)** The helmet, fitted with optically pumped magnetometer sensors (HEDscan, FieldLine Inc.), mounted at the head end of the purpose-built infant bed. **(C)** Positioning of the infant in the OPM-MEG helmet, demonstrated with an infant-sized doll. Infants lay supine and swaddled with the head in the helmet, while high-contrast visual cards were presented above them.

Recordings were performed inside a magnetically shielded room (MSR; MuRoom Small, Cerca Magnetics Ltd, UK; two layers of mu-metal and one layer of copper; internal dimensions 2.4 × 2.4 × 2.4 m; no active shielding), with one caregiver and one experimenter present throughout. Sensors operated in closed-loop mode, measured the magnetic-field component along their local z-axis and were sampled at 5,000 Hz. Infants lay supine and swaddled, naturally awake or asleep, and their position and behaviour were video-recorded throughout from above the bed. During wakefulness, high-contrast visual cards suspended above the helmet, and additional high-contrast visual stimuli when needed, helped to maintain a settled state.

Auditory stimulation began once the caregiver and experimenter agreed that the infant was comfortable and settled and was paused or terminated if the infant became unsettled, required feeding or a nappy change, showed signs of discomfort, or at the caregiver’s request. After the recording, head circumference was measured and, where possible, the infant’s head and helmet were scanned with a handheld 3D scanner (Revopoint POP3; Revopoint 3D Technologies, Shenzhen, China).

### OPM-MEG preprocessing

Data were processed using MATLAB R2019b (MathWorks, Natick, MA, USA), Brainstorm (Tadel et al., 2011), MNE-Python (Gramfort et al., 2013), and procedures adapted from the FLUX pipeline (Ferrante et al., 2022).

Data were downsampled from 5,000 to 1,000 Hz. Continuous recordings were band-pass filtered between 1 and 100 Hz, with notch filters at 50 and 100 Hz to attenuate power-line interference. Signal-space projection (SSP) was used to attenuate cardiac artefacts. Noisy channels were identified by inspecting power spectral density estimates and excluded, and a common set of 45 channels was retained across infants for all analyses. Data were segmented into epochs extending from −1.0 to 2.5 s relative to auditory stimulus onset. Epochs containing high-amplitude artefacts arising from movement, muscle activity, or environmental interference were identified through visual inspection of the OPM-MEG data and of the video recorded during acquisition, and periods of pronounced infant movement identified in the video were rejected.

Infants were presented with 117.7 ± 9.2 happy (range 96-130) and 117.9 ± 9.5 angry trials (range 94-130; t(14) = -0.72, P = 0.49). On average, 20.7 ± 9.8% of happy and 21.4 ± 9.8% of angry trials were rejected per infant (t(14) = -0.76, P = 0.46), leaving 93.5 ± 14.1 happy (range 69–114) and 92.7 ± 14.0 angry trials (range 63–109; t(14) = 0.67, P = 0.51). Trial numbers were then equalized between conditions within each infant using MNE-Python’s equalize_event_counts function (91.2 ± 14.2 trials per condition; range 63-109).

### Event-related field analysis

To compute event-related fields (ERFs), the equalized epochs were decimated to 250 Hz and low-pass filtered at 20 Hz, baseline-corrected using the interval from −210 to −10 ms relative to sentence onset and averaged separately for each condition within each infant. Participant-level ERFs were then averaged to obtain grand-average responses for each condition. Global field power (GFP) computed the standard deviation across the 45 sensors at each time point. Latencies of interest were selected from the condition-pooled grand average without reference to condition: the four global field power (GFP) local maxima at 88, 216, 380 and 1,356 ms.

To compare the timing and amplitude of the evoked response between conditions, each infant’s happy and angry ERFs were projected onto two spatial filters, defined as that infant’s condition-pooled topography at 216 ms and at 380 ms (each averaged over ±10 ms and scaled to unit length). Because the filters were derived from data pooled across conditions with equal numbers of trials, they are independent of the condition contrast. For an early window (140–290 ms; 216-ms filter) and a late window (290–520 ms; 380-ms filter), latency was defined as the time at which the cumulative positive area under the filtered waveform reached 50% (50% fractional-area latency; 4-ms resolution at 250 Hz), and amplitude as the mean within the window.

### Source localization

As individual structural MRI scans could not be acquired, an anatomical template of a 2-month-old infant (O’Reilly et al., 2021), the closest age to the median age of the sample (63 days), was used for source reconstruction in all infants. For the 13 infants with 3D optical scans, the template anatomy was deformed to approximate each infant’s head shape. Because scans were acquired with the infants lying supine, the frontal and lateral surfaces of the head were captured relatively well, whereas the occipital region was not captured in any infant, and the vertex was inadequately captured in most infants. The nasion, left and right preauricular points (LPA, RPA) and four frontal OPM-MEG sensors were identified on each scan in MeshLab (Cignoni et al., 2008).

Using custom Python code implemented with NumPy (Harris et al., 2020), SciPy (Virtanen et al., 2020), and trimesh (Dawson-Haggerty., 2026), the template head surface was aligned with each scan and scaled anteroposteriorly to the nasion, laterally to the preauricular points and vertically to a fixed height-to-length ratio taken from the template and constrained by the frontal and lateral head surfaces captured in the scan, and then adjusted to the measured head circumference without moving the fiducials. The scaled template surface was largely retained where the scan covered the head (median distance to the scan points 1.6 mm), whereas the missing vertex and occipital surfaces were displaced to meet the scan boundary without a step. The resulting head-surface points (150 or 300 per infant) were used to deform the template with Brainstorm’s “Deform default anatomy to fit these points” function.

The sensor array, defined in the helmet coordinate system, was aligned with the scan by a least-squares rigid-body transformation (Kabsch, 1976) fitted to the four frontal sensor positions (root-mean-square residual: median of 0.50 mm, range 0.23–0.63 mm) and coregistered with the deformed template using the nasion, LPA, and RPA (median alignment error 1.0–1.5 mm). For the two infants without scans, whose recording sessions had to be terminated early, the reconstructed head surface of another infant with the same head circumference was used, and coregistration was approximated manually in Brainstorm from photographs taken after recording and video footage captured during acquisition.

Source activity was estimated with weighted minimum-norm estimation (wMNE; Hämäläinen and Ilmoniemi, 1994; Hauk, 2004; Lin et al., 2006) in Brainstorm with the overlapping-spheres method (Huang et al., 1999) computed on each infant’s deformed anatomy, a cortical surface with 15,002 vertices as the source space, and unconstrained source orientations. The noise covariance was estimated from the single-trial baseline period (−210 to −10 ms) and regularised by a factor of 0.1; the inverse solution used a signal-to-noise ratio of 3 and depth weighting with an order of 0.5 (maximum 10). Kernel was computed for the 45 analysed channels. Source amplitude was computed at each cortical vertex as the norm across the three orientations. Because all cortical surfaces were deformed from the same template, vertices corresponded across infants, and source maps were averaged vertex by vertex. At each latency of interest, source amplitude of the condition-pooled ERF was averaged over ±10.

### Time-resolved multivariate pattern analysis

Time-resolved multivariate pattern analysis (MVPA) was performed using MNE-Python (Gramfort et al., 2013) and scikit-learn (Pedregosa et al., 2011) to assess whether neural activity patterns distinguished happy from angry speech over time. Analyses were conducted separately for each infant using preprocessed single-trial sensor-level data (45 channels; equalized trials), decimated from 1,000 to 250 Hz with no additional baseline correction. The analysis interval extended from −0.3 to 1.2 s relative to the prosodically prominent word onset (primary analyses) and, separately, to the sentence onset (secondary analyses).

#### Features

The primary analysis used time-domain waveforms in the 1–20 Hz range broadband; frequency-specific analyses used waveforms and induced power in the delta (1–4 Hz), theta (4–6 Hz), and infant alpha (6–9 Hz) band. Given the 1 Hz high-pass filtering applied during preprocessing, the 1–20 Hz and 1–4 Hz waveforms were obtained by applying additional low-pass filters at 20 and 4 Hz, and 4–6 Hz and 6–9 Hz waveforms by band-pass filtering; all waveform features were single trial signals. To assess whether oscillatory power retained information after removal of the average evoked response, induced power was computed after subtracting the condition-specific mean ERF (estimated from all retained trials) from each trial: Morlet wavelet power (two cycles for 1–4 Hz, three cycles for 4–6 Hz and 6–9 Hz bands) was evaluated at three equally spaced centre frequencies per band (including the lower and excluding the upper boundary), natural-log transformed and averaged within each band, without baseline normalisation. Filtering and power estimation were performed on the full available epochs before extracting the analysis interval for each temporal alignment.

#### Classifier and cross-validation

Spatiotemporal features were constructed using a 12-samples (48-ms) sliding window advanced in 4-ms steps; within each window, values across sensors and samples were concatenated into a single feature vector, and each decoding estimate was assigned to the window midpoint. A logistic regression classifier with the liblinear solver was fitted at each window using MNE-Python’s SlidingEstimator, with feature standardization (StandardScaler) estimated exclusively from the training data in each fold. To ensure that decoding generalized to recordings withheld from training rather than exploiting recording-specific acoustics, we used condition-balanced, utterance-grouped 10-fold cross-validation (Fig. 3). The ten recordings in each condition were shuffled (random seed 42) and assigned to folds with all repetitions of a recording kept together, so that each fold trained on nine happy and nine angry recordings and tested on the remaining one of each. Performance was quantified using the area under the receiver operating characteristic curve (ROC-AUC; chance level 0.5) and averaged across folds.

**Figure 3.**
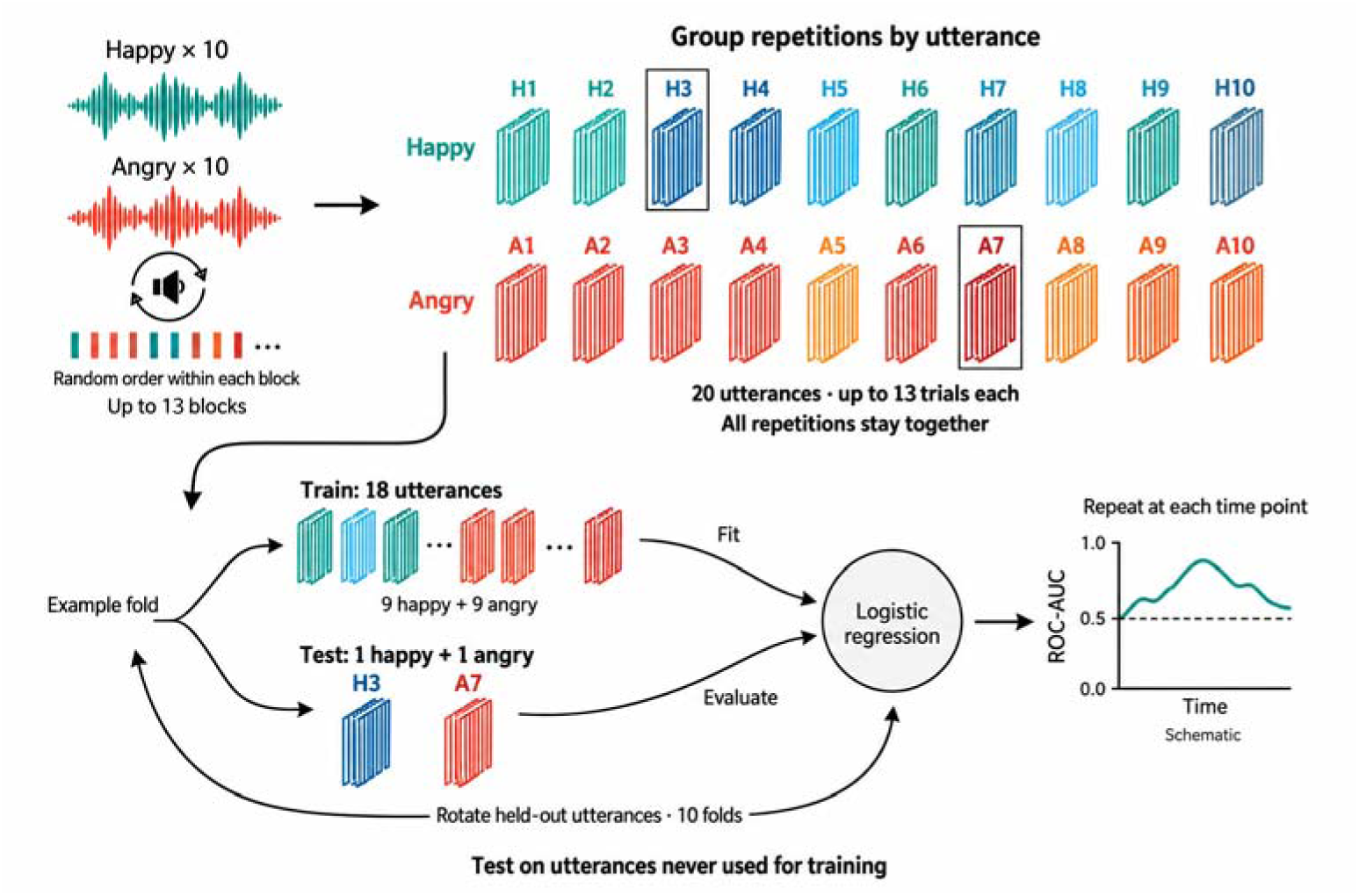
Utterance-grouped 10-fold cross-validation. The stimulus set contains 10 happy (H1–H10) and 10 angry (A1–A10) recordings, each presented up to 13 times. All repetitions of a recording are kept together. The example fold tests H3 and A7 and trains on the remaining 18 utterances; held-out utterances rotate across 10 folds. The illustrated ROC-AUC trace is schematic.

#### Temporal generalization

For the broadband analysis, classifiers trained at each time point were tested at all time points (MNE-Python’s GeneralizingEstimator; same features and folds; time points sampled every 16 ms).

#### Activation patterns

Classifier weights were transformed into activation patterns (A = Cov(X)w; Haufe et al., 2014) using a classifier with the same pipeline, fitted to all trials with the 45 sensors at each time sample. Patterns were normalised by each infant’s maximum absolute value and averaged over the significant cluster window; positive values indicate a larger signal (or power) for happy than for angry speech. For time-domain analyses, cortical patterns were obtained by projecting the window-averaged sensor pattern through each infant’s wMNE kernel. Because log power cannot be projected linearly, cortical patterns for induced power were computed as A(v) = Cov(P_v, ŷ)/Var(ŷ), where P_v is the single-trial induced log power at vertex v (kernel applied to the complex Morlet coefficients of the evoked-subtracted data) and ŷ is the classifier output. Cortical pattern magnitudes were normalised by each infant’s maximum before averaging.

#### Searchlight decoding

Sensor searchlight decoding used each sensor and its six nearest neighbours (seven sensors; 84 features per window). For the broadband analysis, decoding was also performed in 361 overlapping cortical patches; vertices within 15 mm of the centre, at least 10 per patches), using as features the first seven principal components of the patch source signals, computed from the training trials of each fold and stacked over the 12-sample window (84 features, matching the sensor searchlight). Searchlight and patch AUCs were sampled every 16 ms and averaged over each significant cluster window.

#### Acoustic decoding

To compare the timing of neural discrimination with the acoustic information in the stimuli, the same procedure was applied to combined acoustic features (spectral, pitch and intensity) of the 20 recordings, computed on the same 250-Hz grid for both alignments. Decoding was restricted to windows lying entirely within all recordings (58–1,010 ms after sentence onset; 58–950 ms after prominent-word onset). Because each recording contributed a single sample, performance was evaluated with leave-one-pair-out cross-validation over all 100 happy–angry pairs. Significance was assessed with a cluster-based permutation test on recording labels. Acoustic features showed a tendency for above-chance discrimination of emotion early in the utterance, at 58–134 ms after sentence onset (peak AUC = 0.94 at 102 ms) and at 58–150 ms after prominent-word onset (peak AUC = 0.93 at 106 ms), but neither cluster was statistically significant (P = 0.053 for both alignments; Supplementary Fig. 1).

### Statistical analysis

Statistical analyses were conducted using Python with NumPy (Harris et al., 2020), SciPy (Virtanen et al., 2020), MNE-Python (Gramfort et al., 2013) and scikit-learn (Pedregosa et al., 2011). Descriptive statistics are reported as mean ± SD. Unless stated otherwise, tests were two-tailed with α = 0.05, and effect sizes for paired comparisons are reported as Cohen’s d_z.

#### Stimulus and trial comparisons

Acoustic and linguistic properties of the happy and angry recordings (Table 2) and the timing of the temporal landmarks were compared with independent-samples Student’s t-tests (df = 18), with recordings as the unit of analysis. The numbers of trials presented, the numbers of trials retained, and the proportions of trials rejected were compared between conditions across infants with paired t-tests (df = 14).

#### Evoked responses

Time points at which the condition-pooled response differed from baseline (grey shading in Fig. 4A) were identified with one-sample t-tests across infants at each sensor and time point from 0 to 2.0 s, with Benjamini–Hochberg false discovery rate (FDR) correction across all 45 sensors × 501 time points (q < 0.05; Benjamini and Hochberg, 1995). GFP was compared with each infant’s mean baseline GFP using one-tailed t-tests at each time point, FDR-corrected across time points. Latencies and amplitudes of the spatially filtered responses were compared between conditions with paired t-tests, and the phase × condition interaction was tested with a paired t-test on happy − angry differences between the two windows. GFP time courses were compared between conditions with a cluster-based permutation test (0–2.0 s; 2,000 permutations), and mean baseline GFP was compared between conditions with a paired *t*-test. Source amplitude around each latency of interest (Fig. 4C) was compared with the baseline mean using one-tailed spatial cluster-based permutation tests (2,000 permutations).

**Figure 4.**
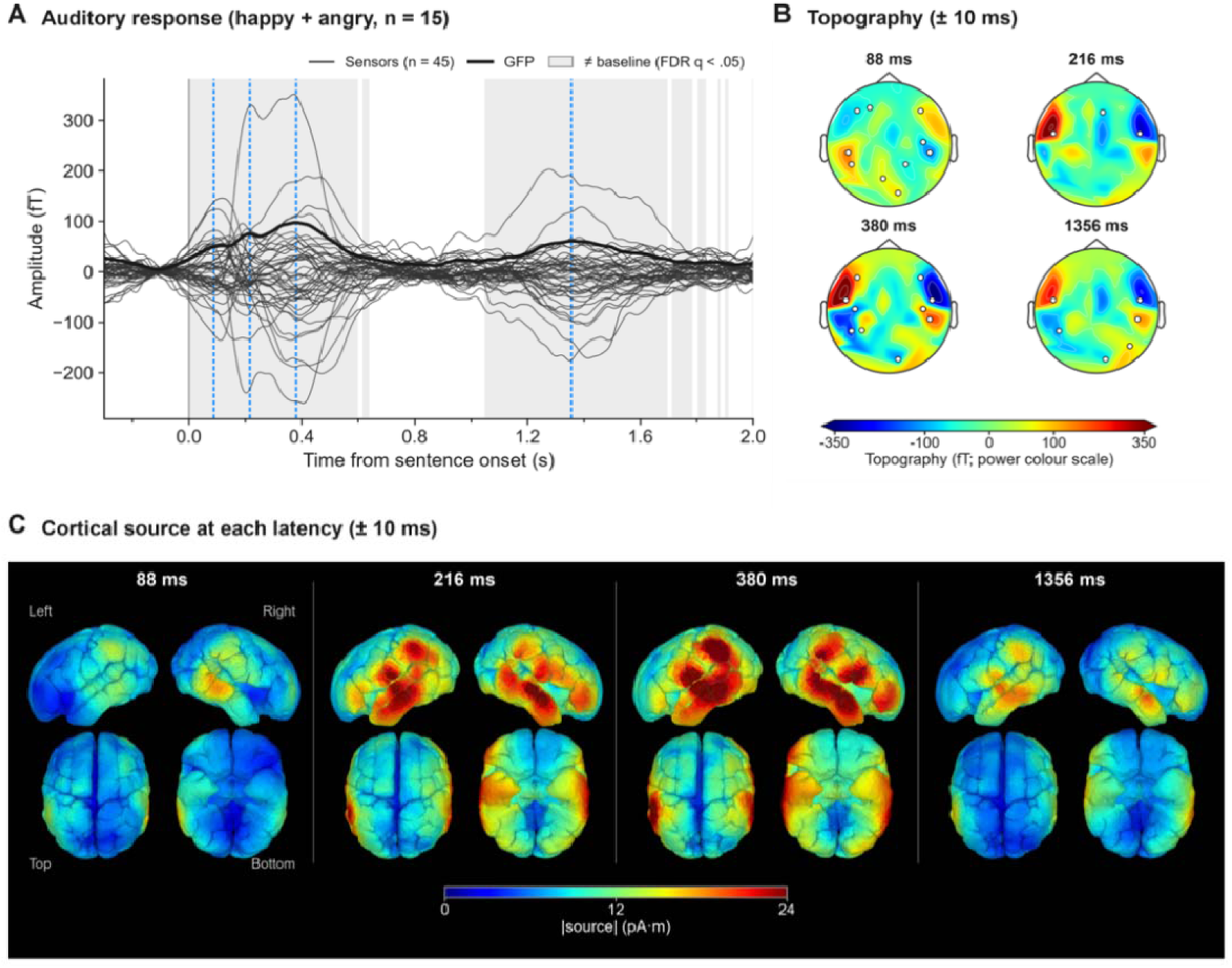
Auditory evoked responses to spoken sentences. **(A)** Auditory response. Grand-average event-related field (ERF) to all sentences (happy and angry pooled; *n* = 15 infants). Thin lines, individual sensors (*n* = 45); thick line, GFP. Grey shading marks time points at which at least one sensor differed from baseline (−210 to −10 ms; two-sided one-sample *t*-tests, FDR *q* < 0.05 across 45 sensors × 501 time points). Blue dashed lines mark the four most prominent GFP peaks (88, 216, 380 and 1,356 ms). **(B)** Topography. Field maps averaged over ±10 ms around each latency. White dots, sensors differing from baseline within this window (FDR *q* < 0.05). **(C)** Cortical sources. Source amplitude averaged over ±10 ms around each latency and across infants, estimated using weighted minimum-norm estimation (wMNE).

#### Decoding

Group-level significance of the decoding time courses was assessed across the 15 infants with a one-tailed cluster-based permutation one-sample *t*-test on participant-level ROC-AUC scores after subtracting 0.5; one-tailed tests were used because only above-chance decoding is interpretable. Adjacent time points exceeding a cluster-forming threshold corresponding to a one-tailed *P* value of 0.05 were grouped into clusters whose statistic was the sum of their *t*-values, and a null distribution of maximum cluster statistics was generated using 2,000 participant-level sign-flip permutations (random seed 42); clusters with a cluster-level *P* value below 0.05 were considered significant. Tests were performed separately for each of the seven analyses and two temporal alignments, correcting for multiple comparisons across time within each analysis; for each analysis, the cluster with the smallest *P* value is reported. Broadband decoding aligned to prosodically prominent word onset was treated as the primary analysis. Across the six frequency-specific analyses (band-limited waveforms and induced power in the delta, theta and alpha bands), cluster-level P values were corrected with the Benjamini–Hochberg FDR procedure within each alignment (Benjamini and Hochberg, 1995), which remains valid under the positive dependence expected among these analyses (Benjamini and Yekutieli, 2001); Uncorrected P values and FDR-adjusted q values are reported. For acoustic decoding, significance was assessed by permuting emotion labels across the 20 recordings (225 permutations, preserving ten recordings per condition), with the same cluster-based procedure applied to the leave-one-pair-out AUC time courses.

## Results

### Infant auditory responses to speech

Spoken sentences elicited a robust auditory response in 2–3-month-old infants (n = 15; happy and angry pooled; Fig. 4A). The response differed from baseline from 0 to 640 ms and from 1,048 to 1,696 ms after sentence onset, with brief further intervals up to 2.0 s (FDR q < 0.05). Global field power (GFP) exceeded its baseline level throughout 0–2.0 s. GFP showed a small peak at 88 ms and prominent peaks at 216, 380 and 1,356 ms.

At 88 ms, the field showed a dipolar pattern over left and right temporal sensors, and from 216 ms onwards, left and right temporal sensors showed the reversed dipolar pattern (Fig. 4B). Differences from baseline were also observed over frontotemporal and parietooccipital sensors.

In the source localization analysis (Fig. 4C), the neural response exhibited distinct spatiotemporal profiles across four peak latencies. At 88 ms, the greatest source amplitude was localized to the right middle temporal cortex. Subsequently, at 216 ms and 380 ms, the signal grew stronger, with the maximum amplitude shifting to the bilateral middle and superior temporal cortex, extending dorsally into the supramarginal cortex and demonstrating significant recruitment of the prefrontal area. At the late latency of 1,356 ms, the most prominent signal was again observed in the bilateral superior and middle temporal cortex. Because source amplitude significantly exceeded the baseline across the entire cortex at all four latencies (spatial cluster P < 0.001), these source maps are interpreted descriptively. Notably, the 1,356-ms peak fell within the range of utterance offsets (1.04–1.60 s).

### Decoding of happy versus angry prosody

We next asked whether happy and angry prosody elicited distinguishable patterns of neural activity. Decoding was time-locked to the onset of the prosodically prominent word (Fig. 1A), and classifiers were always tested on recordings withheld from training (Fig. 3).

Broadband (1–20 Hz) activity discriminated happy from angry prosody 346–454 ms after prominent-word onset (cluster-based permutation test, P = 0.0155; Fig. 5A). Mean AUC within this window was 0.540 ± 0.040 and significantly exceeded the chance level. With sentence-onset alignment, broadband decoding was significant at 418–514 ms (P = 0.0145; mean AUC 0.541, Supplementary Fig. 2). Because the prominent word began within 100 ms of sentence onset, the two alignments captured largely the same effect.In the temporal generalisation matrix, decoding above 0.5 was confined to the diagonal around this window (Fig. 5B). Classifiers trained at one time point did not generalise to distant time points, indicating a transient rather than sustained neural code.

**Figure 5.**
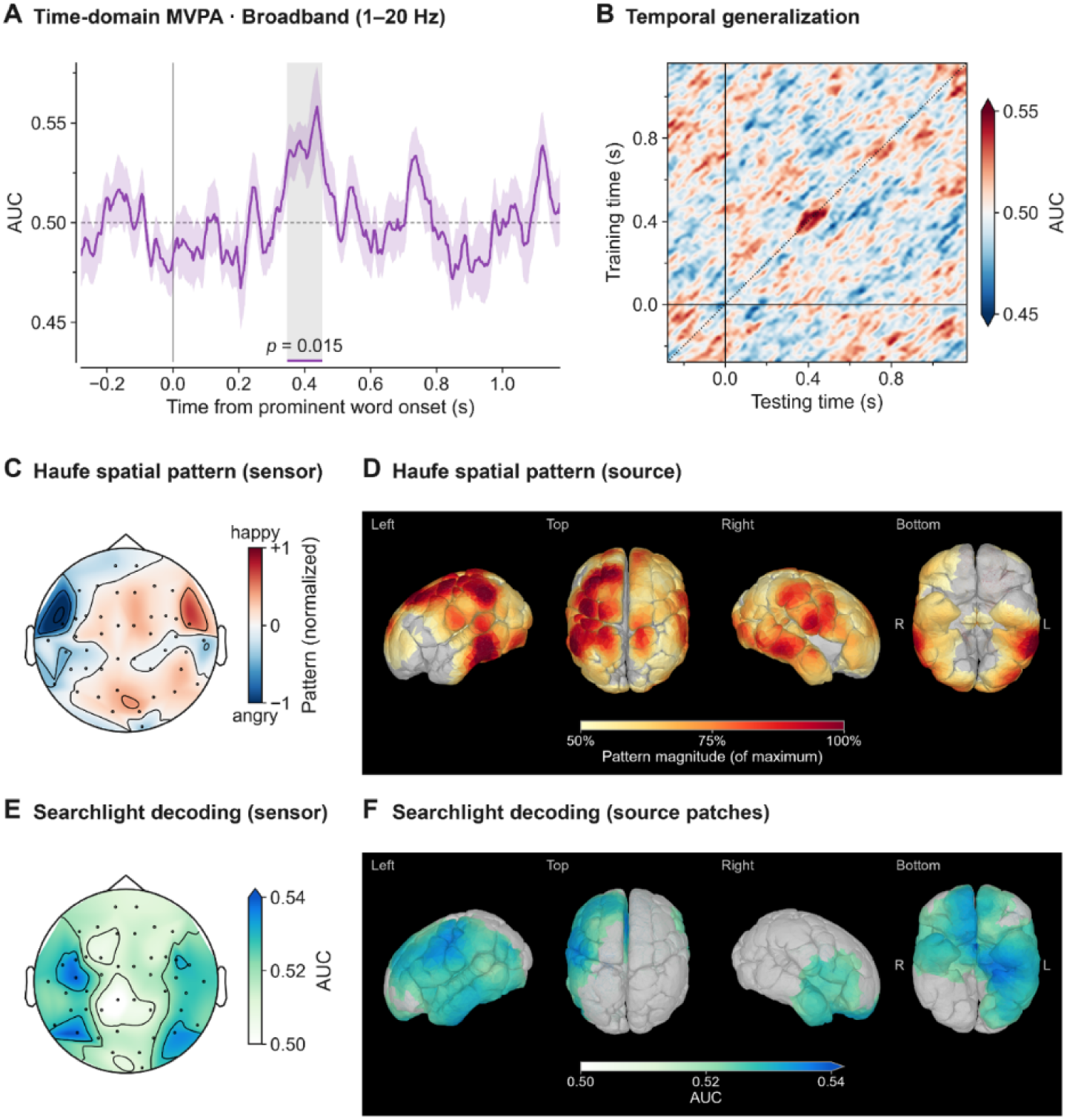
Neural activity discriminates happy from angry prosody after the prosodically prominent word. **(A)** Time-domain decoding. Decoding performance (ROC-AUC) for happy versus angry prosody from broadband (1–20 Hz) signals, time-locked to the onset of the prosodically prominent word. Line, mean across infants (*n* = 15); shading, ± s.e.m.; dashed line, chance (AUC = 0.5). Grey shading and the horizontal bar mark the significant cluster (346–454 ms; one-tailed cluster-based permutation test, *P* = 0.0155). **(B)** Temporal generalisation. Classifiers trained at each time point (y axis) were tested at every time point (x axis); the dotted diagonal corresponds to A. **(C)** Haufe sensor activation pattern. Haufe-transformed classifier weights averaged over the significant window and normalised to [−1, 1], representing interpretable neural sources where red indicates a larger signal for happy and blue indicates a larger signal for angry prosody. Dots, sensor positions. **(D)** Haufe source activation pattern. Magnitude of the source-projected Haufe activation pattern showing the spatial distribution of discriminating neural activity, displayed from 50% to 100% of its maximum. **(E)** Sensor searchlight. Decoding AUC in the significant window for each sensor and its six nearest neighbours, evaluating localized decoding power. **(F)** Source searchlight. Decoding AUC for 361 overlapping cortical patches (radius 15 mm) in the significant window, shown at each vertex as the mean over the patches containing it, identifying informative localized cortical structures; values below AUC = 0.52 are shown in grey.

To identify the specific neural features driving this discrimination, we computed the Haufe activation pattern, which transforms classifier weights into interpretable neural signals. The pattern was strongest over bilateral temporal sensors (Fig. 5C) and resembled the bilateral temporal topography of the evoked response (Fig. 4B), but with opposite polarity. Because the pattern was negative over left temporal and positive over right temporal sensors, where the evoked field had the opposite sign, it indicates larger temporal fields for angry than for happy prosody in this window. In the source-space activation map derived via Haufe transformation, which highlights the neural activity pattern most distinctively underlying the discrimination of happy versus angry prosody within the left hemisphere (fronto-temporo-parietal network), strong activation was distributed across the left precentral cortex and caudal middle frontal cortex (anterosuperior regions), the supramarginal cortex (posterosuperior region), and the entire inferior and middle temporal cortex (inferolateral region), with additional localized patches visible near the right temporal and frontal areas (Fig. 5D).

Searchlight analyses, employed to systematically map the localized distribution of decoding informativeness across the entire brain, converged on similar regions. The most informative sensors lay over the left temporal, left posterior temporal, and right posterior temporal regions (mean AUC 0.53–0.54; Fig. 5E). Correspondingly, the most informative cortical patches were identified within the left postcentral gyrus, inferior parietal lobule, supramarginal cortex, left fusiform gyrus, lateral occipito-temporal cortex, as well as the right orbitofrontal cortex and anterior temporal cortex (Fig. 5F).

### Latency and amplitude of evoked responses

To test whether the patterns discovered in the time-decoded MVPA also manifested as differences in conventional evoked-response measures, we compared sentence-aligned ERFs between conditions. Happy and angry sentences elicited responses with similar topographies and time courses (Fig. 6A), and their GFP time courses did not differ (no happy–angry cluster, smallest cluster P = 0.67; baseline GFP, P = 0.66; Fig. 6B).

**Figure 6.**
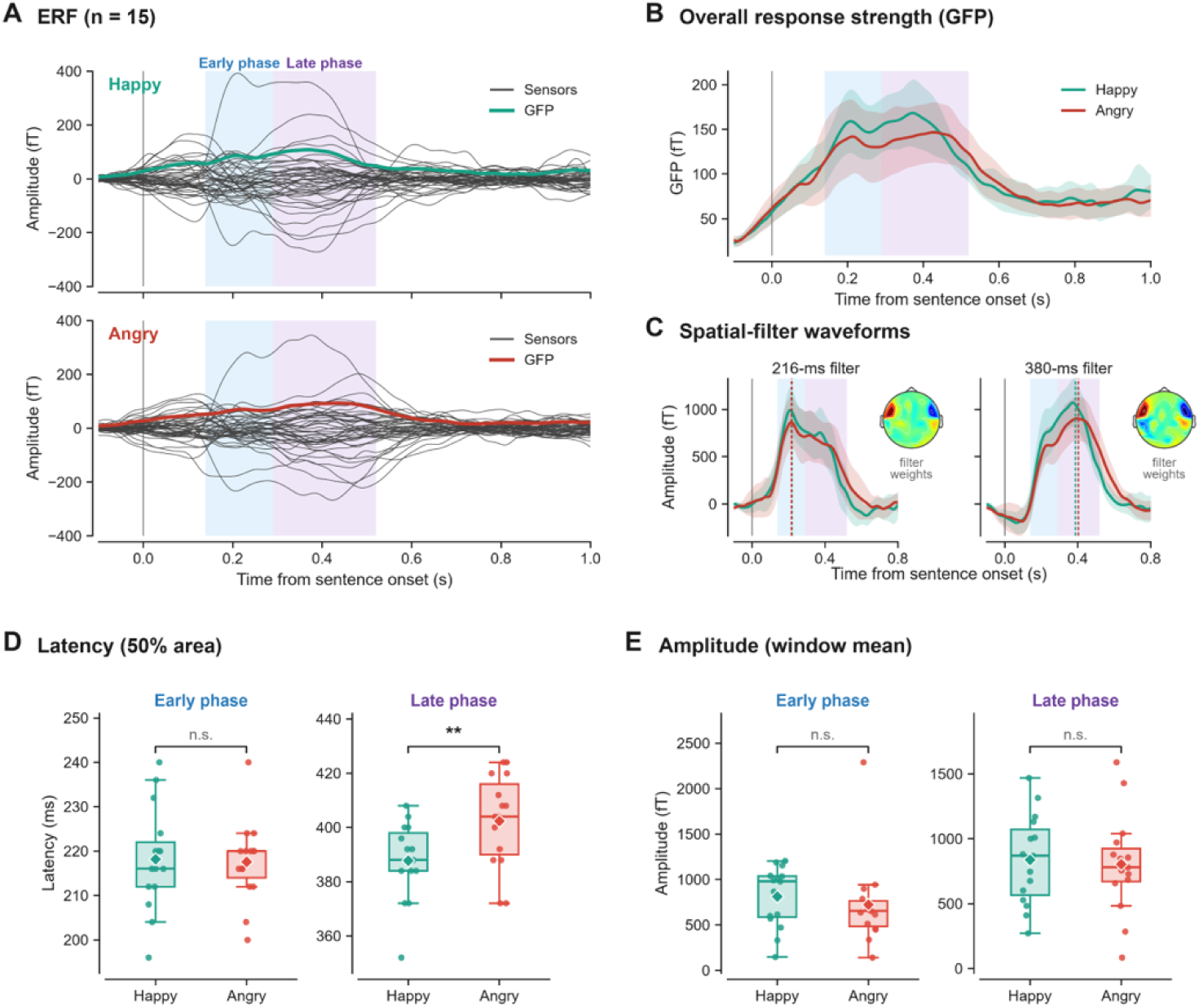
Angry prosody delays the late phase of the auditory response without changing its amplitude. **(A)** ERFs. Grand-average ERFs (*n* = 15) to happy (top) and angry (bottom) sentences, time-locked to sentence onset. Thin lines, sensors; thick line, GFP. Blue and purple shading mark the early (140–290 ms) and late (290–520 ms) phases. **(B)** Overall response strength. GFP for happy (green) and angry (red) sentences; shading, 95% CI. **(C)** Spatial-filter waveforms. Each infant’s ERF projected onto spatial filters defined from that infant’s condition-pooled topographies at 216 ms (left) and 380 ms (right); insets show filter weights (cf. Fig. 4B). Dashed lines, peak latency per condition; shading, 95% CI. **(D)** Latency. 50% fractional-area latency of the early (216-ms filter) and late (380-ms filter) phases, showing a significant condition difference exclusively in the late phase (*t*(14) = −3.73, *P* = 0.002). **(E)** Amplitude. Mean amplitude within the same windows, demonstrating no significant differences between happy and angry prosody in either the early or late phase. Box plots show the median and interquartile range; whiskers, 1.5 × IQR; diamonds, means; dots, individual infants. ** *P* < 0.01; n.s., not significant (two-sided paired *t*-tests).

We quantified the timing and amplitude of the two main response phases using spatial filters derived from each infant’s condition-pooled topographies at 216 and 380 ms, which are therefore unbiased with respect to the happy–angry contrast (Fig. 6C). Early-phase latency did not differ between prosodies (happy 218.1 ms, angry 217.6 ms; t(14) = 0.23, P = 0.82; Fig. 6D). In contrast, the late phase occurred 14.7 ms earlier for happy than for angry prosody (happy 387.7 ms, angry 402.4 ms; t(14) = −3.73, P = 0.002, d_z = −0.96). The happy–angry latency difference was larger in the late than in the early phase (interaction: t(14) = −3.73, P = 0.002). This late-phase delay for angry prosody persisted when responses were aligned to the onset of the prominent word (−14.9 ms, t(14) = −4.56, P < 0.001). Amplitude did not differ in either phase (early: t(14) = 0.74, P = 0.47; late: t(14) = 0.36, P = 0.73; Fig. 6E).

Thus, emotional prosody modulated the timing of the late auditory response but not its amplitude.

### Frequency-specific decoding

To further identify the frequency components of the discriminative signal, we performed decoding in the delta (1–4 Hz), theta (4–6 Hz), and infant alpha (6–9 Hz) bands using both band-limited waveforms and induced power (Fig. 7).

**Figure 7.**
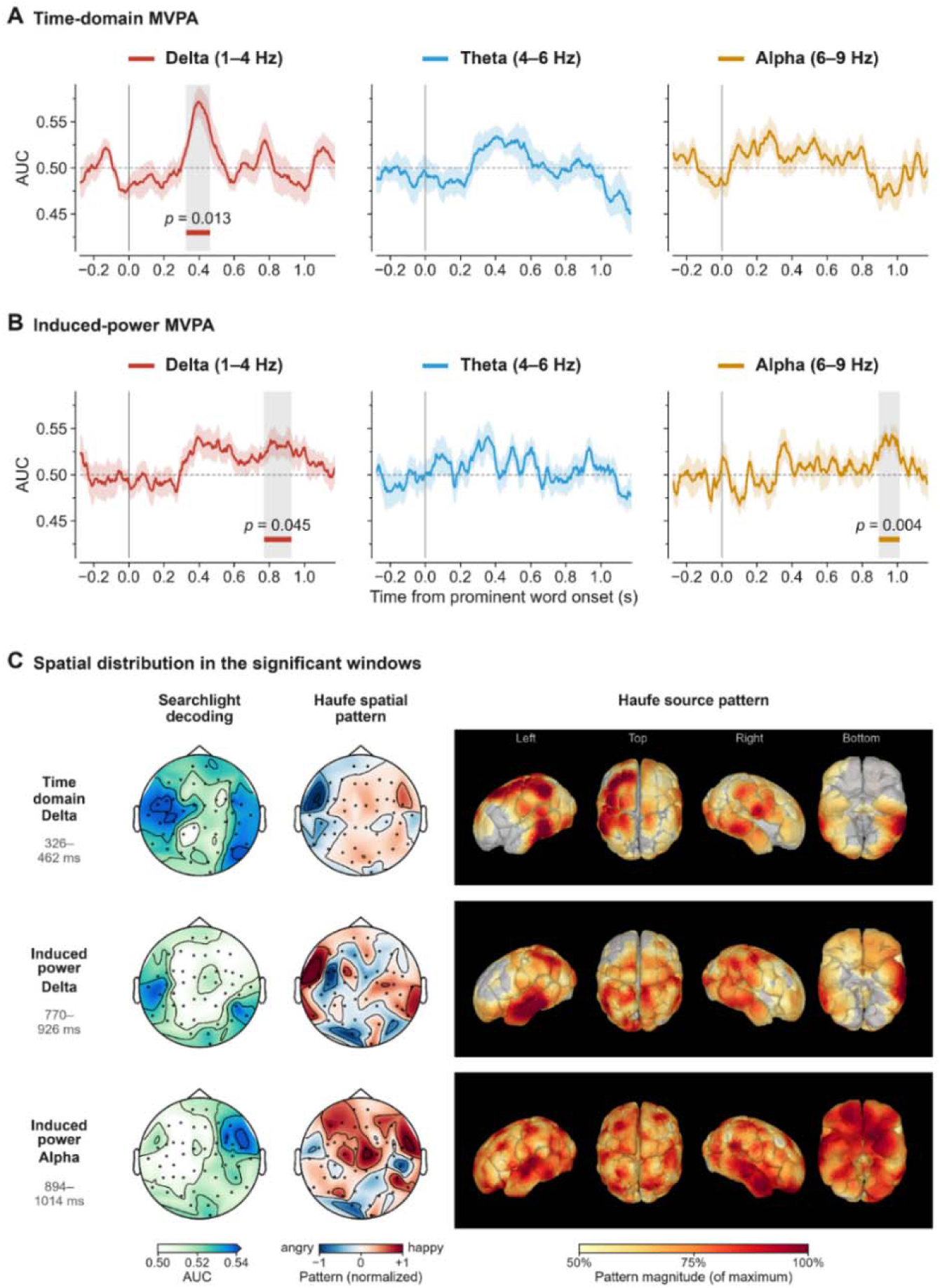
Frequency-resolved decoding of emotional prosody. **(A)** Time-domain decoding. Decoding based on band-limited waveforms in the delta (1–4 Hz), theta (4–6 Hz), and infant alpha (6–9 Hz) bands, time-locked to the onset of the prominent word. **(B)** Induced power decoding. Decoding based on log-transformed Morlet wavelet power, calculated after subtracting the average ERF of each condition from single trials. In A and B, solid lines represent the mean across infants (*n* = 15), shaded areas represent the standard error (\(\pm \) s.e.m.), and grey shaded intervals and coloured bars indicate significant clusters (one-tailed cluster-based permutation test). *P*-values are cluster-level and uncorrected across analyses. FDR-adjusted *q* values are reported in the main text. **(C)** Spatial distributions within significant time windows: time-domain delta (326–462 ms), induced delta (770–926 ms), and induced alpha (894–1,014 ms). Left: sensor searchlight AUC; Middle: sensor activation patterns (red indicates larger for happy prosody, blue indicates larger for angry prosody); Right: source activation pattern magnitudes (50–100% of maximum).

For band-limited waveforms, delta activity discriminated between the two prosodies in the 326–462 ms interval after prominent-word onset (Fig. 7A; P = 0.0135, FDR q = 0.041; mean AUC 0.554 ± 0.050), and the time window, sensor patterns, and source patterns of this effect closely matched the broadband results (Fig. 7C, top row; cf. Fig. 5). Theta decoding rose moderately at 370–510 ms but did not reach significance (P = 0.07, q = 0.105), and alpha decoding showed no significant clusters (P = 0.24, q = 0.288). The six most informative searchlight sensors were strongly localized over the left temporal cortex (AUC 0.54).

Induced power analysis yielded later effects (Fig. 7B). Induced alpha power discriminated the two prosodies in the 894–1,014 ms interval (P = 0.0035, q = 0.021; mean AUC 0.534 ± 0.028), and induced delta power discriminated the prosodies in the 770–926 ms interval (P = 0.045; mean AUC 0.531 ± 0.041) that did not survive FDR correction (q = 0.090). Induced theta power failed to discriminate the two prosodies (P = 0.32, q = 0.32). The Haufe sensor pattern for induced delta was largest at the left temporal and right posterior temporal sensors, and the source pattern was strongest in the left superior/middle temporal cortex and left parietal cortex (Fig. 7C, middle row). Searchlight decoding also showed clusters in the left temporal and right posterior temporal sensors. The Haufe sensor pattern for induced alpha showed predominantly positive weights across most sensors (78% of all sensors), meaning that alpha power was higher for happy prosody than angry prosody in most sensors (Fig. 7C, bottom row). When viewed in the source space, this pattern was broadly distributed without a clear cortical focus. Searchlight decoding results showed the largest clusters in the right temporal and fronto-temporal sensors.

Taken together, the discrimination of emotional prosody emerged first in phase-locked activity (most prominently in the delta band) at 346–454 ms after the prominent word, followed by induced alpha power at approximately 900 ms.

## Discussion

Previous evidence for early neural sensitivity to emotional prosody has come from isolated syllables or haemodynamic measures, leaving open when such sensitivity emerges within natural speech (Cheng et al., 2012; Zhang et al., 2017, 2019; Shekhar et al., 2019). Combining OPM-MEG with time-resolved MVPA, we addressed this question in 2- to 3-month-old infants listening to natural sentences. Sentences elicited auditory responses at about 88, 216, 380 and 1,356 ms after onset. Neural patterns distinguished happy from angry prosody 346–454 ms after the onset of the prosodically prominent word, mainly through phase-locked delta-band activity, although global field power (GFP) did not differ between conditions. Angry prosody also delayed the late auditory response by about 15 ms, and induced alpha power differed between conditions at a later window (894–1,014 ms). These findings show that, by 2–3 months of age, the infant brain distinguishes emotional prosody in natural sentences within few hundred milliseconds of a silent prosodic event. They also demonstrate that OPM-MEG combined with time-resolved decoding can recover information from spatially distributed infant brain responses even when overall response strength does not differ.

### Infant auditory responses to spoken sentences

GFP pooled across all sentences peaked at 88, 216, 380 and 1,356 ms. In this paper, polarity was defined as positive when the source dipole pointed in the superior direction. Following paediatric MEG studies (Paetau et al., 1995; Yoshimura et al., 2014; An et al., 2020), we label the positive response at 216 ms as the P1m. Other studies refer to the same response as the P2m or P2 (Edgar et al., 2015; Chen et al., 2023; Corvilain et al., 2025). In this scheme, the 88 ms response is an early response of opposite polarity that precedes the P1m, and it may correspond to the N1m described at about 100 ms by Chen et al. (2023).

At 88 ms, a dipolar pattern appeared over the left and right temporal sensors, and the source was maximal in the right middle temporal cortex. This suggests an early response originating in the auditory cortex, but infant responses in this time range have rarely been studied. Chen et al. (2023) described an N1m at about 100 ms in most infants but did not analyse it. Studies presenting pure tones to infants younger than 2 months did not report brain activity earlier than about 250 ms (Lutter et al., 2006; Corvilain et al., 2025). Moreover, because the adult-like N1m does not emerge reliably until late childhood (Paetau et al., 1995; Ponton & Eggermont, 2001), this response is unlikely to be the same component as the adult N1m despite the shared label. The 88 ms response can therefore be interpreted as reflecting early acoustic processing of speech onset at this age. This response may have been rarely reported in previous studies because OPM sensors are positioned closer to the scalp than conventional MEG sensors, which yields larger signal amplitude of small early responses, or because of the longer inter-stimulus interval. The source was right-lateralised. Because this response was pooled across all sentences, it cannot be interpreted as an emotion-specific effect, but it is consistent with reports that the processing of acoustic change is right-lateralised in infants (Homae et al., 2006; Telkemeyer et al., 2009).

The 216 ms response showed a bilateral temporal dipolar pattern with polarity opposite to that of the 88 ms response. Its sources were located in the bilateral superior and middle temporal regions, and this component appears to correspond to the P1m reported in the literature. When pure tones were presented to infants younger than 2 months, the P1m occurred at about 250 ms (Lutter et al., 2006; Corvilain et al., 2025) and shortened with age (Lutter et al., 2004). When 500 Hz tones were presented to infants and toddlers aged 2–24 months, it occurred at 111–241 ms, with a mean of about 167–180 ms in infants younger than 12 months (Chen et al., 2023). The 216 ms latency observed here falls between these values, within the range expected for 2- to 3-month-old infants. In particular, a study using speech stimuli reported a P1m at about 200 ms in 2- to 3-month-old infants (Yoshimura et al., 2020); our 216 ms response therefore closely matches the value reported for the same age and the same type of stimulus, and is also consistent with the hypothesis that the main auditory response would occur at around 200 ms. P1m latency shortens rapidly during the first year (Chen et al., 2023). Later in childhood, the P1m remains prominent in response to voice stimuli, and its latency continues to shorten (Yoshimura et al., 2014; An et al., 2020; Kotecha et al., 2009). Longitudinal evidence suggests that the prominent early voice-evoked response observed in infancy may correspond to the P1m component observed in children over 2 years of age (Yoshimura et al., 2020).

The 380 ms response had the same polarity as the 216 ms response and may be a late positive component that is consistently observed in infancy. In infant EEG, the P150–N250–P350– N450 components are present from birth, and their latencies shorten with age (Kushnerenko et al., 2002). In infant MEG, a positive component at about 400 ms (P4m) has also been described among responses that alternate in polarity in the order N1m–P2m–N3m–P4m (Chen et al., 2023). At 380 ms, sources extended from the temporal cortex to supramarginal and prefrontal regions. These regions overlap with those reported to respond emotional prosody in haemodynamic studies. In 2-month-old infants, the left temporoparietal cortex responded more strongly to happy speech (Shekhar et al., 2019), and in newborns the region around the right supramarginal gyrus was sensitive to fearful prosody (Zhang et al., 2017). In adults, this region is involved in phonological processing (Hartwigsen et al., 2010) and in reorienting attention to salient stimuli (Corbetta & Shulman, 2002). Frontal regions also responded differentially to emotional prosody in 7-month-old infants (Grossmann et al., 2010b) and in newborns (Zhang et al., 2019).

At 1,356 ms, we observed the largest peak in the period after 380 ms; this time point fell within the range of utterance offsets (1.04–1.60 s). The maximal source amplitude at this time was again located in the bilateral superior and middle temporal cortex. This response may, in the first instance, be an offset response to the end of the sentence. In adults, the end of a sound elicits an offset response in the same auditory cortical regions as the onset response, with a smaller amplitude than the onset response (Pantev et al., 1996). In infants aged about 2–4 months, clear responses have also been observed at both the onset and the offset of vowel stimuli (Uhler et al., 2018), and newborn ERPs already reflect differences in vowel offset duration (Urbanec et al., 2024). That the late response in the present study appeared in the same temporal regions as the early responses and was smaller than the 380 ms response is consistent with these characteristics. However, because sentence durations varied from 1.04 to 1.60 s, offset responses would be expected to be temporally dispersed in averaged responses time-locked to sentence onset.

### Discrimination of happy and angry prosody

Broadband (1–20 Hz) neural activity significantly discriminated happy from angry prosody at 346–454 ms after prominent-word onset (mean AUC 0.540 ± 0.040). Because this discrimination occurred before the end of the sentence, it suggests that the infant brain distinguished the prosodic difference in real time, before hearing the whole utterance. This time window also overlaps with the late response windows reported in infant emotional prosody studies. In 7-month-old infants, angry prosody elicited a more negative ERP than happy and neutral prosody (Grossmann et al., 2005), and in 3- to 12-month-old infants, differences in mismatch responses between emotional prosodies appeared at 100–200 ms and 300–500 ms (Kao & Zhang, 2025). This is consistent with the prediction that discrimination would emerge in a window several hundred milliseconds after onset, later than the early sensory responses. To the best of our knowledge, this is the first study to show that such discrimination can be detected as time-resolved neural patterns in 2- to 3-month-old infants, and in sentence contexts rather than in isolated words.

In the temporal generalisation matrix, decoding was confined to the vicinity of the diagonal. This suggests that the neural patterns carrying discriminative information constitute a dynamic code that changes rapidly over time (King & Dehaene, 2014). However, in infant data with a short decoding window and a low signal-to-noise ratio, generalisation away from the diagonal may have gone undetected owing to insufficient statistical power, so the absence of a sustained code cannot be concluded. Furthermore, whereas GFP showed no difference between happy and angry prosody, MVPA revealed significant decoding. Because GFP collapses signal strength across all sensors into a single value, it cannot capture differences in spatial patterns. Multivariate analysis, in contrast, is sensitive to differences in spatial patterns across sensors and, because a decoder is trained for each individual, can also exploit spatial patterns that differ between infants (Petit et al., 2020; Ashton et al., 2022). MVPA has also been used to extract phonetic information from speech in the EEG of 3-month-old infants (Gennari et al., 2021).

Angry prosody delayed the latency of the late phase by about 14.7 ms but did not affect the latency of the early phase or the amplitude of either phase. This effect persisted when data were aligned to prominent-word onset (about 14.9 ms). That the difference appeared only at a later stage rather than at initial detection suggests that processing of angry prosody was more prolonged or required additional processing. In previous infant studies, angry prosody was also distinguished from other prosodies in late time windows. The late negative response in 7-month-old infants was interpreted as reflecting increased attentional allocation to angry signals (Grossmann et al., 2005), and in 3- to 12-month-old infants angry prosody elicited a more negative late mismatch response than sad prosody (Kao & Zhang, 2025). However, because these studies reported amplitude differences whereas the present study reports a latency difference, direct comparison is limited.

Conversely, the late phase was faster for happy prosody, which is consistent in direction with a haemodynamic study of infants of the same age. In a study measuring the left hemisphere of 21 2-month-old infants with diffuse optical tomography, happy speech elicited greater responses than angry speech in the left STG and posterior STS, and greater responses than neutral speech in the temporoparietal cortex (Shekhar et al., 2019). Although the measures differ (the magnitude of the haemodynamic response versus response latency), both results point in the same direction, suggesting that happy prosody may have a processing advantage in superior temporal regions at this age. Early sensitivity to positive prosody has also been reported in other studies. Newborns opened their eyes more to happy speech in their native language (Mastropieri & Turkewitz, 1999), and non-nutritive sucking was more suppressed by sad and neutral utterances than by happy utterances (Silvestri et al., 2026). In newborn fNIRS, happy prosody elicited greater activation than negative prosody in the left superior frontal gyrus and angular gyrus (Zhang et al., 2019), and 3- to 12-month-old infants listened longer to happy than to angry prosody (Kao et al., 2022). Six-month-old infants’ preference for infant-directed speech has been explained as a preference for positive emotion (Singh et al., 2002), and the prosody of infant-directed speech can be regarded as the result of freely expressing emotion (Trainor et al., 2000). Greater exposure to speech carrying positive emotion may therefore have contributed to the faster processing of happy prosody.

Nevertheless, this happy advantage should not be generalised. In newborns, fearful syllables elicited larger mismatch responses than happy and neutral syllables (Cheng et al., 2012), and the region around the right supramarginal gyrus was more sensitive to fearful prosody (Zhang et al., 2017). In a study of facial expressions in 5- to 12-month-old infants, the N290 was larger for fearful and happy faces, whereas the P400 and Nc were larger for angry faces (Xie et al., 2019). In other words, the effects of emotional valence depend on the measure and the processing stage. A negativity bias has been reported mainly in studies from the second half of the first year onwards, particularly in social referencing studies (Vaish et al., 2008). Our results therefore show only a temporal advantage for happy prosody in the late auditory response of 2- to 3-month-old infants, and do not imply a general positivity bias.

### Contributions of frequency bands

In the present study, the broadband decoding effect was driven mainly by the delta band (1–4 Hz). Delta band-limited waveforms significantly discriminated happy from angry prosody at 326–462 ms, and their time window and sensor and source patterns were nearly identical to the broadband results. In the theta band, AUC increased somewhat at 370–510 ms but not significantly, and no significant cluster was found in the alpha band. This indicates that the discriminative information was carried mainly by phase-locked activity in the delta band, that is, by low-frequency evoked responses. Because infant auditory evoked responses are dominated by slow components with periods of several hundred milliseconds, this result can be regarded as the late-phase latency difference described above being expressed mainly in the delta band. The delta band has been proposed as a suitable timescale for processing slow prosodic structure at the sentence and phrase level (Giraud & Poeppel, 2012; Goswami, 2011). The infant brain also tracks the speech envelope in the delta and theta bands from 4 months of age, and infant-directed speech has particularly strong amplitude modulation at around 2 Hz (Attaheri et al., 2022; Leong et al., 2017). Newborns can already discriminate languages on the basis of rhythmic information (Nazzi et al., 1998). In this light, our results suggest that such low-frequency processing may also be involved in the discrimination of emotional prosody.

In induced power, alpha power discriminated the two prosodies at a later time window, 894– 1,014 ms, with higher power for happy prosody. Suppression of alpha-band power has been used as an index of increased attention, and in 6-month-old infants, power in the infant alpha band (6–9 Hz) is suppressed during sustained attention (Xie et al., 2018). Applying this interpretation, the relatively lower alpha power for angry prosody may reflect more sustained attention to angry prosody. This is consistent with the delayed late phase for angry prosody in the present study, and with the interpretation of 7-month-old infants’ responses to angry prosody as reflecting increased attentional allocation (Grossmann et al., 2005).

### Spatial distribution of discriminative information

We interpreted the spatial information of the decoder using two measures. Haufe activation patterns show the spatial distribution and direction of the signals used by a decoder trained on all sensors (Haufe et al., 2014), whereas searchlight analysis shows where the two prosodies can be discriminated from local signals alone. At the sensor level, Haufe patterns showed opposite polarities over the left and right temporal regions, matching the magnetic field distribution of bilateral auditory cortical responses. The decoder therefore most likely exploited differences in bilateral auditory cortical responses, which is also consistent with the late-phase latency effect observed in the same time window. The difference in sign between hemispheres arises from the geometry of the magnetic field and is not considered to indicate hemisphere-specific emotional processing.

In contrast, searchlight analysis showed that discriminative information was lateralised to the left hemisphere. At the sensor level, left temporal and posterior temporal sensors were the most informative. At the source level, information was broadly distributed, centred on the left temporal cortex and extending to the lateral frontoparietal convexity (precentral and postcentral gyri, caudal middle frontal gyrus, supramarginal region and inferior parietal lobule) and the ventral temporo-occipital surface, whereas in the right hemisphere it appeared only weakly in orbitofrontal and anterior temporal regions. Haufe source patterns also highlighted similar left-hemisphere regions but were more widely distributed. That the Haufe patterns were bilateral whereas the searchlight showed left lateralisation indicates that, although the decoder used responses from both hemispheres, the information discriminable from local signals alone was greater on the left.

This left-hemisphere lateralisation is in line with reports that the presence of emotion and the discrimination of emotional valence in prosody are processed in different hemispheres. In newborns, the right superior temporal gyrus responded more strongly to emotional than to neutral prosody, whereas differences between happy and negative prosody appeared in the left superior frontal gyrus and angular gyrus (Zhang et al., 2019). In 2-month-old infants, the left STG and posterior STS also responded more strongly to happy than to angry speech (Shekhar et al., 2019). Because the present study compared happy and angry prosody, which differ in valence, the concentration of discriminative information in the left hemisphere is consistent with these findings.

The contribution of the lateral frontoparietal convexity should be interpreted with caution. Given reports that Broca’s area is activated when 3-month-old infants listen to sentences (Dehaene-Lambertz et al., 2006) and the view that the sensorimotor system may contribute to speech perception even before infants begin to produce speech (Choi et al., 2023), sensorimotor and frontal regions may have been involved. However, these regions may reflect left temporal information being co-localised to adjacent cortex, or factors such as attentional shifts or movement (Corbetta & Shulman, 2002). The weak contribution of right orbitofrontal and anterior temporal regions is partly consistent with reports that emotional vocalisations modulate orbitofrontal responses in 3- to 7-month-old infants (Blasi et al., 2011).

### Limitations and future directions

Several limitations should be considered. First, to obtain a sufficient number of trials within a short recording time, we compared only two emotions. Happy and angry prosody differ in valence but are both relatively high in arousal. We therefore cannot determine whether the observed neural differences reflect valence, specific emotion categories, or the accompanying acoustic differences (fundamental frequency, intensity and speech rate). In addition, because there was no neutral condition, we could not separate responses to the presence of emotional prosody (Zhang et al., 2019) from differences between valences. Second, as is common in infant neuroimaging studies, the sample was not large (n = 15) and decoding accuracy was low. Third, because infants of this age sleep for long periods and data collection is difficult, recordings included a mixture of sleep and awake states. Because infants have relatively shallow sleep, the difference may not be as large as in adults; nevertheless, sleep state may still have altered auditory responses and responses to emotional prosody. Future studies should follow larger samples longitudinally to examine how emotional prosody processing develops over the first year of life. Because OPM sensor arrays can be adapted to head size, they allow measurements from infancy to adulthood (Hill et al., 2019), and recordings are possible even in 1-month-old infants (Corvilain et al., 2025). Such longitudinal studies can therefore now be conducted from early infancy. Furthermore, including neutral prosody and emotions that differ in arousal would make it possible to separate the effects of the presence of emotion, valence and arousal.

## Conclusion

Using OPM-MEG with time-resolved decoding, we showed that the brains of 2- to 3-month-old infants distinguish happy from angry prosody in natural sentences within a few hundred milliseconds of a prosodically prominent word, mainly through delta-band evoked activity centred on the left temporal cortex. Notably, this temporal information was not evident in global field power but emerged from the spatial patterns exploited by the decoder. These findings indicate that neural sensitivity to emotional prosody, previously shown with isolated syllables and haemodynamic measures, already operates on the timescale of natural speech in early infancy. More broadly, they show that combining OPM-MEG recordings, which pair millisecond resolution with rich spatial information, with multivariate decoding can reveal when information is present in the infant brain even when it is not expressed in overall response strength, which is difficult to achieve with EEG or haemodynamic measures alone. This opens the way for longitudinal studies of how emotional prosody processing develops across the first year of life.

## Acknowledgments

We are especially grateful to the infants and their parents who participated in this study. We thank Jiaqi Li for advice on the multivariate pattern analysis. This work was supported by a UKRI BBSRC Pioneer Award (BB/Y51309X/1) to K.A., by start-up funding and LES equipment bid from the University of Birmingham to K.A., by a Wellcome Trust Discovery Award (227420) to O.J., by the NIHR Oxford Health Biomedical Research Centre (NIHR203316) to O.J., by a UKRI EPSRC Quantum Technologies Career Development Fellowship (EP/W028050/1) to A.K., and by a UKRI ESRC Research Grant (UKRI1889) to A.B. The funders had no role in study design, data collection and analysis, the decision to publish, or preparation of the manuscript.

## Author Contributions

J.Y. Data curation; Formal analysis; Methodology; Software; Visualization; Writing – original draft; Writing – review & editing. V.M.L. Data curation; Formal analysis; Methodology; Visualization; Writing – original draft; Writing – review & editing. A.W. Investigation; Writing – review & editing. A.K. Funding acquisition; Investigation; Writing – review & editing. A.B. Conceptualization; Funding acquisition; Supervision; Writing – review & editing. B.P. Investigation; Writing – review & editing. G.O. Investigation; Writing – review & editing. O.J. Conceptualization; Funding acquisition; Methodology; Supervision; Writing – review & editing. K.A. Conceptualization; Data curation; Formal analysis; Funding acquisition; Investigation; Methodology; Project administration; Supervision; Visualization; Writing – original draft; Writing – review & editing.

## Conflicts of interests

The authors declare no competing interests.

## Data availability statement

The data supporting the findings of this study are available from the corresponding author upon reasonable request, subject to ethical and institutional requirements. The data are not publicly available because sharing of paediatric neuroimaging data is restricted by privacy and ethical considerations. The analysis code is available from the corresponding author upon reasonable request.

## Supplementary Figures

**Supplementary Figure 1.**
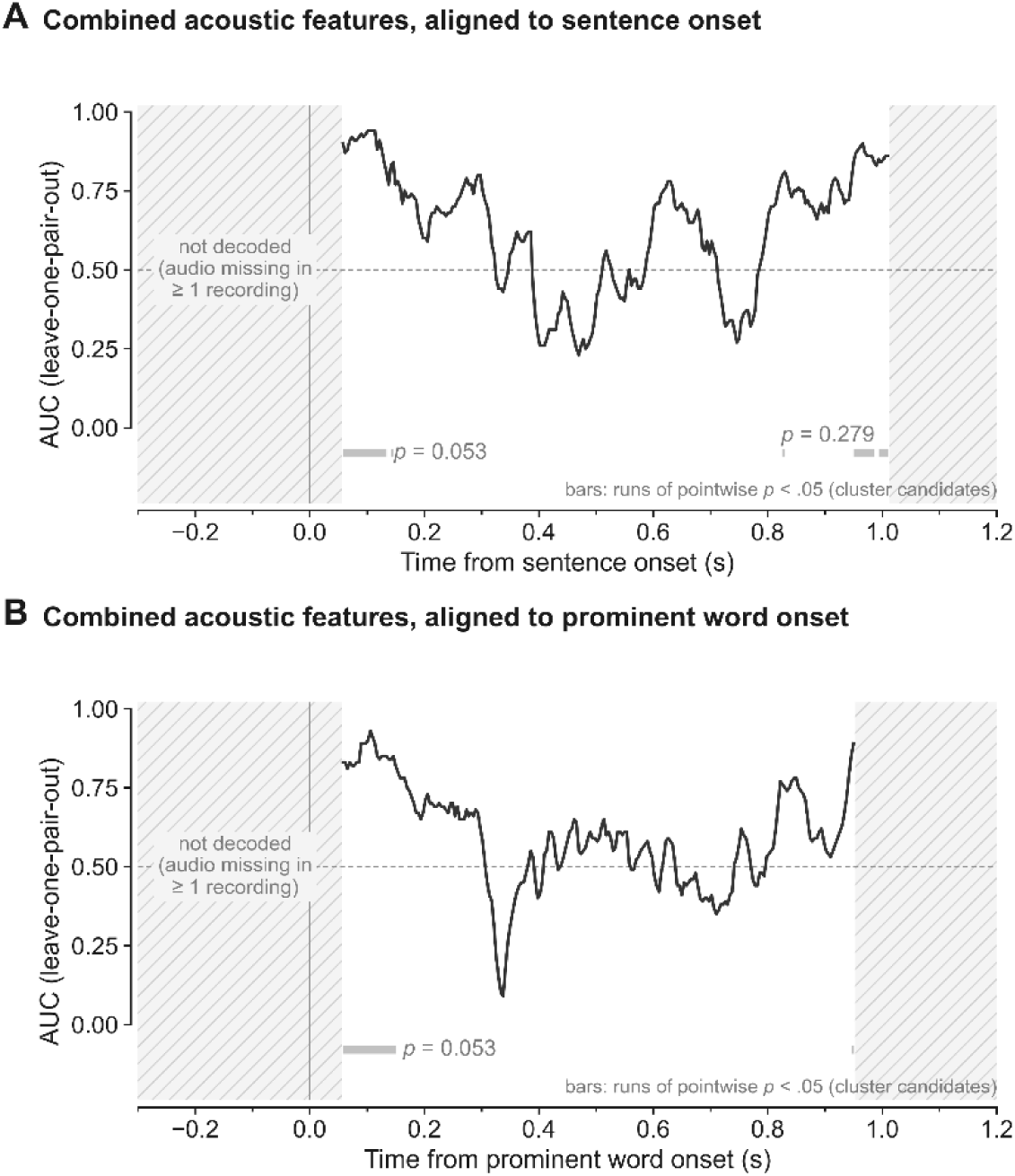
Acoustic decoding

**Supplementary Figure 2.**
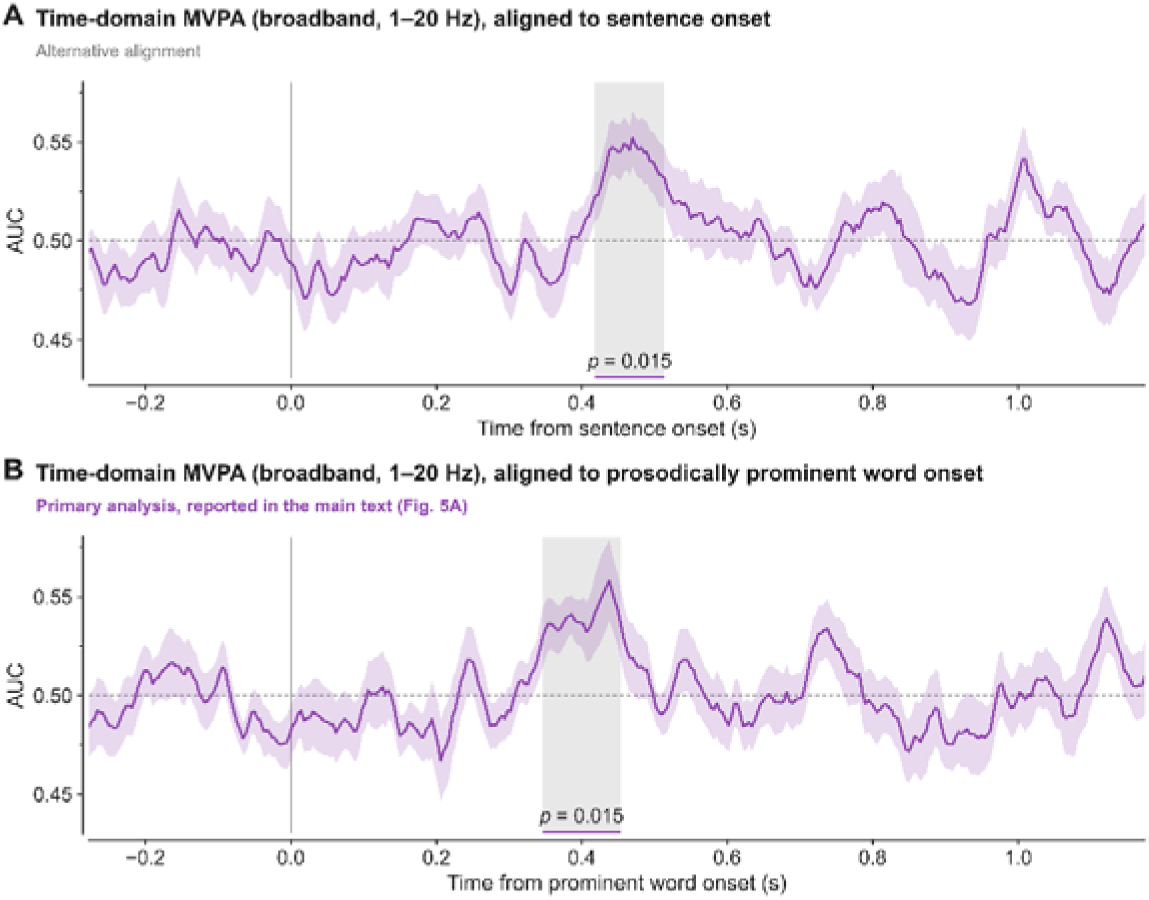
MVPA analysis compared between the sentence onset and prosodically prominent word onset

## Notes

### Competing Interest Statement

The authors have declared no competing interest.

